# 123FASTQ: an intuitive and efficient tool for preprocessing Illumina FASTQ reads

**DOI:** 10.1101/2024.03.08.584032

**Authors:** Milad Eidi, Samaneh Abdolalizadeh, Mohammad Hossein Nasirpour, Javad Zahiri, Masoud Garshasbi

## Abstract

Next-generation sequencing (NGS) has revolutionized genetic research, but preprocessing raw sequencing reads remains a crucial step. Existing tools for quality control (QC) and trimming have limitations, leading to fragmented workflows. In this article, we present 123FASTQ, a comprehensive and user-centric software solution that integrates QC analysis, trimming, and format conversion in a userfriendly interface. 123FASTQ combines the advantages of existing tools while addressing their limitations. It offers improved QC analysis with insightful visualizations, semi-automated trimming recommendations based on QC results, and various trimming options. The software also supports different file formats and provides efficient performance. By simplifying NGS data preprocessing, 123FASTQ enables researchers to utilize high-quality reads for downstream analyses. It is cross-platform and freely available at https://sourceforge.net/projects/project-123ngs/.

## Introduction

Next-generation sequencing (NGS) has revolutionized the field of genetics, enabling researchers to analyze DNA and RNA sequences at an unprecedented scale and speed. A crucial step in the analysis of NGS data is the preprocessing of raw sequencing reads, which involves quality control (QC) and trimming procedures. Various tools have been developed to address these preprocessing tasks, each with its own advantages and limitations. In this article, we present 123FASTQ, a comprehensive and user-centric software solution designed to overcome the shortcomings of existing NGS read preprocessing tools. FastQC and its emulation, Falco are widely used tools for QC analysis of NGS data(1,2). These provide a superior graphical user interface (GUI) and command line interface (CLI) for quality evaluation, offering visualizations and metrics to assess the quality of raw sequencing reads. However, these are primarily focus on quality assessment and does not support additional preprocessing tasks such as trimming or format conversion. This limitation necessitates the use of additional tools to perform these tasks, leading to a fragmented workflow for researchers. Trimmomatic and CutAdapt are powerful tools known for their robust trimming functions(3,4). These tools offer a wide range of trimming options, including adapter removal, quality-based trimming, and length filtering. However, it lacks built-in QC analysis capabilities and relies heavily on command-line interfaces, which can be challenging for users with limited bioinformatics experience. Another popular tool, Fastp, integrates QC analysis and trimming functions(5). It provides a command line interface and supports various QC metrics for assessing reads quality. However, Fastp may not cater to the diverse needs of researchers, as it does not provide comprehensive and informative QC plots. In contrast to the aforementioned tools, 123FASTQ offers a unified solution by integrating essential preprocessing tasks within an easy-to-navigate GUI. It combines the advantages of FastQC’s quality assessment, Trimmomatic’s trimming capabilities, and additional features such as format conversion. By providing a user-friendly interface, 123FASTQ aims to make Illumina NGS read preprocessing more accessible and efficient for researchers with varying levels of bioinformatics expertise. Furthermore, 123FASTQ enhances the preprocessing workflow by incorporating semi-automated trimming recommendations based on QC results. This feature empowers users to trim their data quickly while considering the software’s recommendations, streamlining the decision-making process. Overall, 123FASTQ addresses the limitations of existing preprocessing tools by offering a comprehensive suite of features, user-friendly interface, and compatibility with different file formats. By amalgamating the finest features of existing tools, 123FASTQ strives to simplify NGS data preprocessing, ensuring the utilization of high-quality reads for downstream analyses.

## Materials and Methods

### Software Development

123FASTQ was developed as a tool for preprocessing Illumina NGS reads. It was implemented using the Java programming language, ensuring compatibility across different operating systems. 123FASTQ does not require any specific hardware configuration, however performances are dependent on the number of CPU cores and the amount of available memory (See result section).

### Quality Control (QC) Analysis

123FASTQ incorporates basic codes from the FASTQC tool for performing QC analysis of raw sequencing reads(1). However, numerous improvements were made to enhance the usefulness and graphical appeal of the QC plots. In addition, we integrated an interactive bubble chart that visually represents the distribution of mean qualities corresponding to various read lengths. The size of each bubble correlates with the abundance of reads possessing specific quality and length attributes, providing an insightful overview. QC analysis can be performed in both single-mode and comparative mode, allowing users to compare results between different read files or sets of read files. Input files can be in FASTQ, bz2 and gz compressed FASTQ, FAST5, SAM, and BAM formats.

### Semi-Automated Trimming

One of the key innovations of 123FASTQ is the provision of trim recommendations based on the QC results of input FASTQ files. When users submit single-end or paired-end FASTQ files for QC analysis, a button appears in the result panel. Clicking on this button opens a trimming tab and pre-fills trimming options based on the QC results. This semi-automated trimming approach enables users to perform trimming quickly and also allows them to apply their own trimming strategies while considering 123FASTQ recommendations.

### Trimming Process

123FASTQ incorporates various trimming options to empower users to clean reads effectively within a visual environment. Some of the features from Trimmomatic, such as the adapter decontaminator, MAXINFO, LEADING, and TRAILING, are embedded in 123FASTQ(4). The software accepts input files in FASTQ or compressed FASTQ formats (bz2, gz, or zip). For other formats like SAM, BAM, and FAST5, users are recommended to use the 123FASTQ converters to convert them to the FASTQ format before performing trimming.

### Format Conversion

123FASTQ includes two format converters in its converters panel, allowing users to convert Fast5 and SAM/BAM formats to FASTQ format for further processing.

## Results

As 123FASTQ is capable of managing different tasks such as QC analysis, trimming, and format conversion, it is crucial to assess its performance in each of these aspects when compared to other widely recognized tools. To guarantee a fair evaluation, we carried out the benchmark on a computer with identical specifications and utilized the same FASTQ files. Our assessment of the benchmarked tools focused on factors like time and memory usage for the QC task, and time, memory usage, and alignment accuracy for the trimming task.

In the QC analysis, 123FASTQ showed similar performance to FASTQC, but our tool stands out with improved quality and more insightful results. When it comes to trimming, our tool demonstrated a processing speed comparable to Trimmomatic, with a slight edge in flexibility.

## Discussion

Next-generation sequencing (NGS) has brought about a transformative change in the genomics domain, empowering scientists to examine DNA and RNA sequences at an unparalleled magnitude and velocity. A crucial initial step in the analysis of NGS data is the preprocessing of raw sequencing reads, encompassing quality control (QC) and trimming procedures. This stage is vital for securing the precision and dependability of subsequent analyses. While various software tools have been devised for preprocessing NGS reads, the majority concentrate on particular aspects of the process and might not accommodate the requirements of all users, particularly those with minimal bioinformatics background. In this article, we present 123FASTQ, an all-inclusive and user-centric software solution designed to overcome the shortcomings of existing NGS read preprocessing tools. Our software embeds essential features such as QC and trimming within an easy-to-navigate graphical user interface (GUI) that caters to users with diverse levels of expertise.

Current preprocessing tools, including FastQC, Trimmomatic, CutAdapt, and Fastp, exhibit their own advantages and drawbacks. For example, FastQC provides a superior GUI for quality evaluation but does not support additional preprocessing tasks, while Trimmomatic and CutAdapt boast robust trimming functions but lack QC and format conversion capabilities. Moreover, these tools predominantly rely on command-line interfaces, which can pose difficulties for inexperienced users. 123FASTQ resolves these issues by consolidating all necessary preprocessing tasks into a single, user-friendly GUI. Our software not only simplifies the preprocessing workflow but also ensures that only the highest quality reads are utilized for downstream analyses. By amalgamating the finest features of existing tools and presenting a user-friendly interface, 123FASTQ strives to render NGS data preprocessing more accessible and efficient for researchers across all skill levels.

Additionally, 123FASTQ offers a comprehensive suite of features, making it a one-stop solution for NGS read preprocessing. The software supports various file formats and can seamlessly handle both single-end and paired-end sequencing data. With its advanced quality assessment and visualization capabilities, users can easily identify and address issues in their data. Furthermore, 123FASTQ provides a range of trimming and filtering options, including adapter removal, quality-based trimming, and length filtering, ensuring that the data meets the desired quality standards. The software also offers parallel processing and optimization for improved efficiency, enabling researchers to quickly process large datasets. By providing a versatile and adaptable solution, 123FASTQ equips scientists with a powerful tool to streamline their NGS data preprocessing and achieve reliable, high-quality results.

## Notes

https://sourceforge.net/projects/project-123ngs/

### Competing Interest Statement

The authors have declared no competing interest.

